# Comparative Analysis and Data Provenance for 1,113 Bacterial Genome Assemblies

**DOI:** 10.1101/2021.12.14.472616

**Authors:** David A. Yarmosh, Juan G. Lopera, Nikhita P. Puthuveetil, Patrick Ford Combs, Amy L. Reese, Corina Tabron, Amanda E. Pierola, James Duncan, Samuel R. Greenfield, Robert Marlow, Stephen King, Marco A. Riojas, John Bagnoli, Briana Benton, Jonathan L. Jacobs

**Author notes:** EDAN Diagnostics; Madison, WI (USA).

## Abstract

The quality and traceability of microbial genomics data in public databases is deteriorating as they rapidly expand and struggle to cope with data curation challenges. While the availability of public genomic data has become essential for modern life sciences research, the curation of the data is a growing area of concern that has significant real-world impacts on public health epidemiology, drug discovery, and environmental biosurveillance research^1–6^. While public microbial genome databases such as NCBI’s RefSeq database leverage the scalability of crowd sourcing for growth, they do not require data provenance to the original biological source materials or accurate descriptions of how the data was produced^7^. Here, we describe the *de novo* assembly of 1,113 bacterial genome references produced from authenticated materials sourced from the American Type Culture Collection (ATCC), each with full data provenance. Over 98% of these ATCC Standard Reference Genomes (ASRGs) are superior to assemblies for comparable strains found in NCBI’s RefSeq database. Comparative genomics analysis revealed significant issues in RefSeq bacterial genome assemblies related to genome completeness, mutations, structural differences, metadata errors, and gaps in traceability to the original biological source materials. For example, nearly half of RefSeq assemblies lack details on sample source information, sequencing technology, or bioinformatics methods. We suggest there is an intrinsic connection between the quality of genomic metadata, the traceability of the data, and the methods used to produce them with the quality of the resulting genome assemblies themselves. Our results highlight common problems with “ reference genomes” and underscore the importance of data provenance for precision science and reproducibility. These gaps in metadata accuracy and data provenance represent an “ elephant in the room” for microbial genomics research, but addressing these issues would require raising the level of accountability for data depositors and our own expectations of data quality.

---

The National Center for Biotechnology Information’s (NCBI) RefSeq database has become an essential cornerstone of the global genomics research community, but declining data quality and the increasing cost of manual data curation by end-users are growing areas of concern ^8,1,3,2^. As RefSeq continues to expand, so too does the risk for data errors, omission, obfuscation, or falsification to go undetected and to potentially damage trust in this enormously important public resource ^9,10^. RefSeq contains over 357,657 prokaryotic genome assemblies. It is the largest collection of non-redundant, annotated genome assemblies available, and it is built exclusively from crowd-sourced data. However, despite extensive efforts to create automated curation pipelines and tools to improve RefSeq data, significant quality issues remain in genome assemblies found within RefSeq ^11–13^. For example, while all newly deposited prokaryote genome assemblies are automatically annotated, the associated metadata records (i.e., BioSample, BioProject, SRA, Assembly data) are submitted by depositors who are not required to provide attribution for the biological materials behind each genome ^14,15^. In fact, the International Nucleotide Sequence Database Collaboration (INSDC) policy states “*the quality and accuracy of the record are the responsibility of the submitting author, not of the database*,” which is to say that metadata, which are often crucial for comparative genomics research, are not curated or verified for accuracy ^16^. This is further complicated by data omissions, lack of controlled vocabulary for terms, variable taxonomic naming conventions, and competing metadata package formats. In many cases, tracing the provenance of an individual assembly to its source material in order to verify its authenticity becomes challenging, and manual curation is frequently required to detect and correct RefSeq metadata errors ^17^.

In this study, we present the results of an ongoing whole-genome sequencing initiative at the American Type Culture Collection (ATCC) to provide end-to-end data provenance from source materials to reference-grade microbial genomes, hereafter referred to as “ATCC Standard Reference Genomes” (ASRGs). Presented here are 1,113 bacterial ASRGs from authenticated materials that were produced via a hybrid *de novo* assembly approach. We compared them to assemblies in RefSeq where metadata indicated they were produced by 3^rd^ party labs from materials sourced from ATCC For 366 ASRGs (∼33%), we were able use metadata to compare them to one or more assemblies in RefSeq. The remaining 747 ASRGs (∼66%) represented potentially novel assemblies. All ASRGs described here are available for research use via the ATCC Genome Portal (https://genomes.atcc.org)^18^.

## Whole-genome Sequencing of 1,113 ATCC Bacterial Strains

High-molecular-weight genomic DNA (HMW-gDNA) was extracted from 1,113 bacterial strains obtained from ATCC’s biorepository. Each strain was cultured using strain-specific protocols and subjected to quality control (QC) for contamination, viability, purity, phenotype, and taxonomic identity (Figure 1). For whole-genome sequencing (WGS), HMW-gDNA was split and subjected to sequencing using both Illumina and Oxford Nanopore Technologies (ONT) next-generation sequencing (NGS) platforms (Figure 1). Next, reads were taxonomically classified using One Codex’s metagenomics platform to assess the purity of each NGS library prior to *de novo* assembly ^19^. Read sets were then down-sampled to predetermined coverage depths (Illumina, 100x; ONT, 60x) expected to be optimal for bacterial genome assemblies ^20–22^. Lastly, a hybrid-assembly pipeline incorporating reads from both platforms produced *de novo* assemblies for each strain ^23^. High-level summary metrics for each ASRG are shown in Table S1 and Fig. 2. All 1,113 ASRG assemblies were estimated to be over 95% complete by *CheckM*; 1,015 were found to be over 99% complete and 329 are 100% complete ^24^. A total of 617 are considered high-quality, closed genome references.

**Fig 1.**
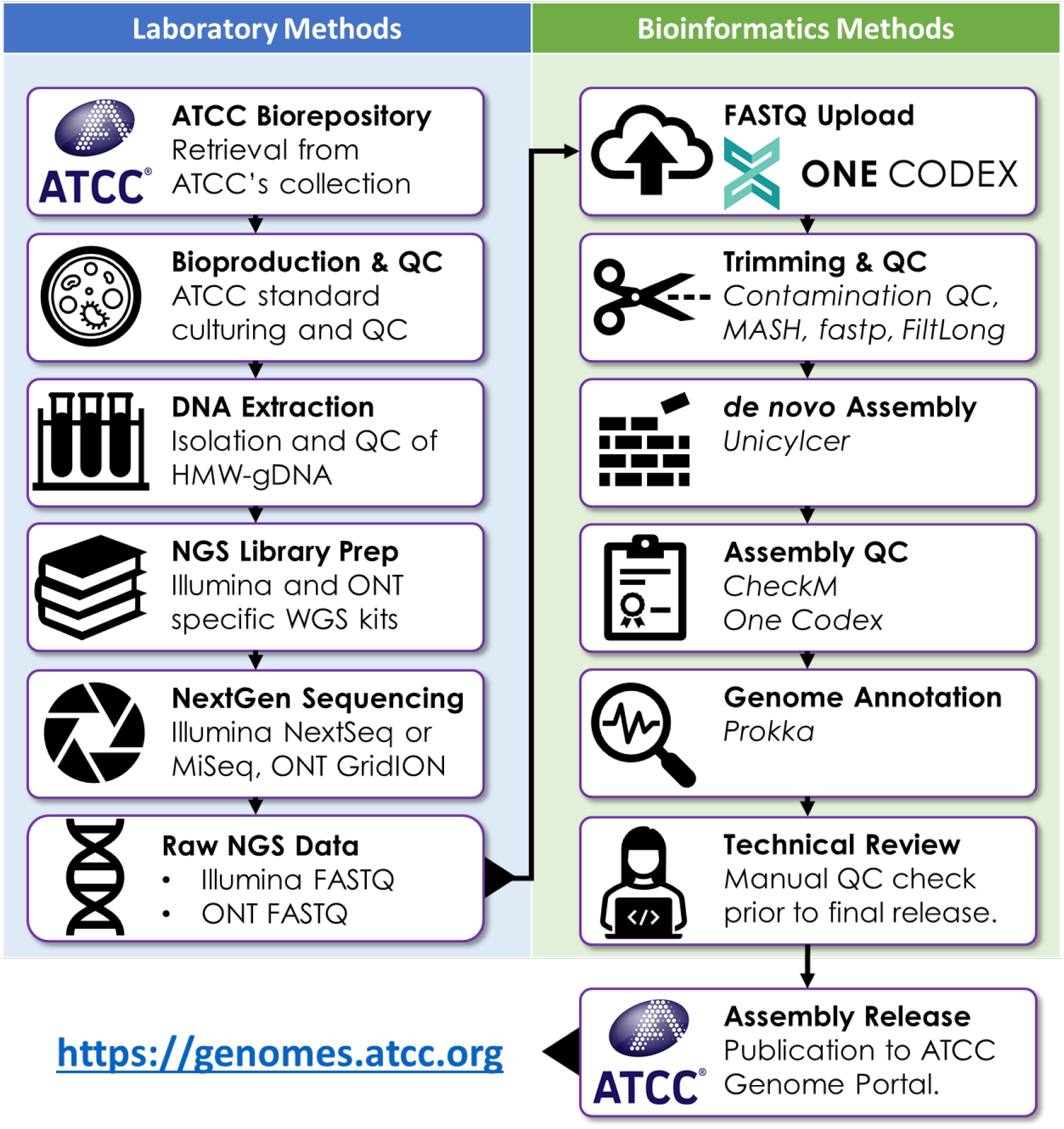
A Pipeline for End-to-End Genomic Data Provenance. Source materials are obtained directly from the ATCC biorepository and tracked through to the final assembly and genome annotation. Upfront culture conditions varied depending on the species cultured, but downstream process steps were performed using standardized protocols for DNA extraction, library prep, sequencing, and bioinformatics. Each pipeline is hosted on One Codex’s cloud infrastructure.

**Fig 2.**
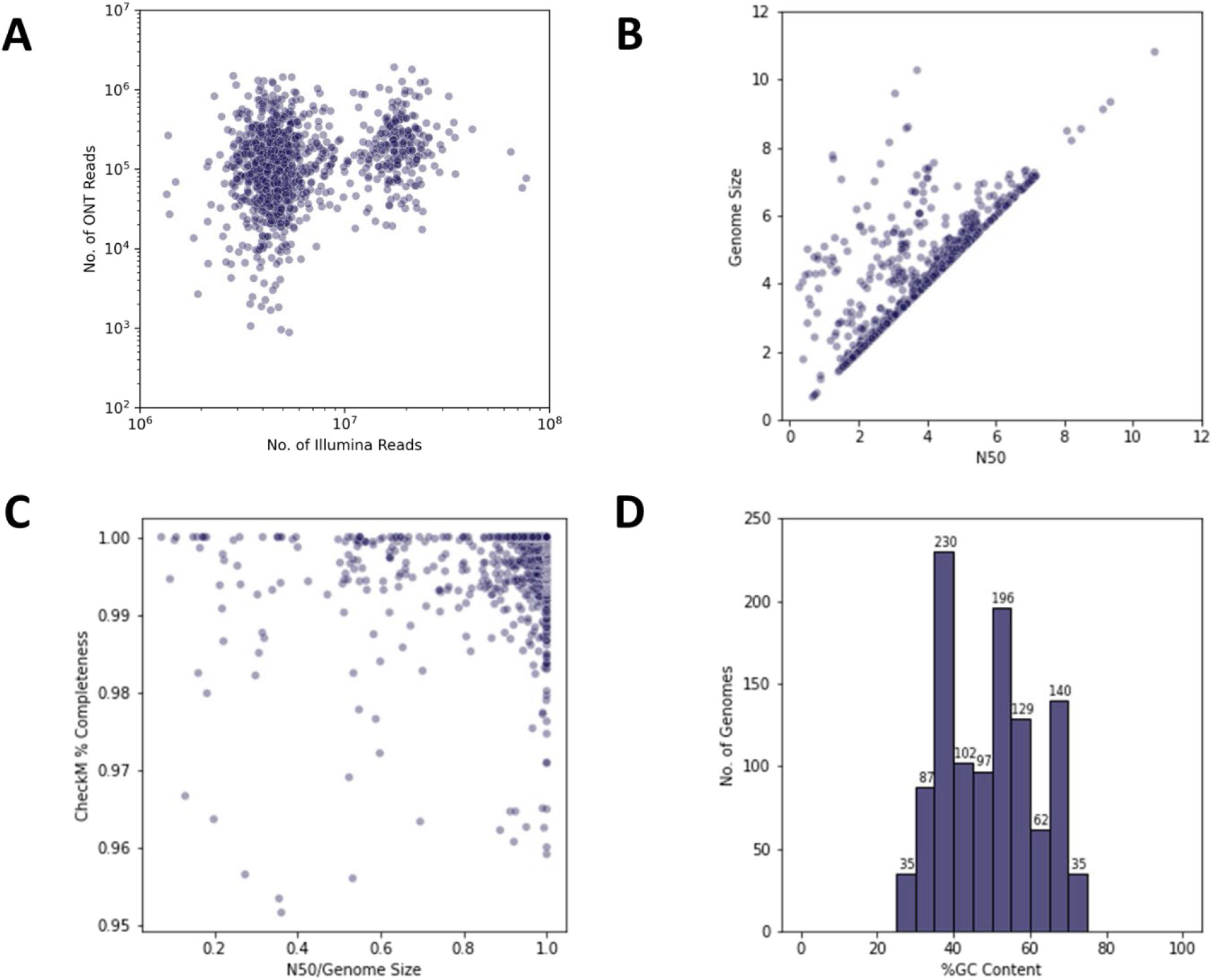
Sequencing & Quality Metrics for 1,113 Bacterial Genome Assemblies. **(A)** Illumina vs. ONT reads for ASRGs before down-sampling. **(B)** N50 metrics vs. genome size. **(C)** N50 normalized by genome size vs *CheckM* genome completion estimates. **(D)** Diversity of G:C% content for all 1,113 ASRG assemblies.

## Survey of Bacterial Genome Assemblies in RefSeq

We compared the ASRG assemblies to those in NCBI’s RefSeq Bacteria Database (https://ftp.ncbi.nlm.nih.gov/genomes/refseq/bacteria/) labeled as representing ATCC bacterial strains, i.e., assemblies where the ATCC strain name (or a synonymous name) was indicated in the title, description, or other metadata field in the GenBank assembly record. We intentionally did not search RefSeq using a traditional comparative genomics approach (i.e., by sequence homology, BLAST, etc.) since this would require arbitrary thresholds for determining strain identity, and metadata descriptors are intended to be useful for these types of queries. Using this approach, we found 2,701 genome assemblies in RefSeq, which collectively comprised 1,960 different ATCC strains (Table S2, Fig. 3A). Interestingly, RefSeq had numerous examples of bacterial strains represented by multiple assemblies or submitted by different groups, and it often included “strains” resulting from intentional genetic modification (i.e., there are 33 different RefSeq assemblies for *Serratia marcescens* subsp. *marcescens* ATCC^®^ 13880™). This is despite it representing a “non-redundant” database. Overall, we found one or more duplicate assemblies in RefSeq for 158 strains for which we also produced an ASRG, including instances of assemblies for genetically modified strains mislabeled as representing “type strains” (See Table S2). These errors and strain duplications create risks for researchers who may unwittingly use these data in their own research yet remain unaware of these issues.

**Fig 3.**
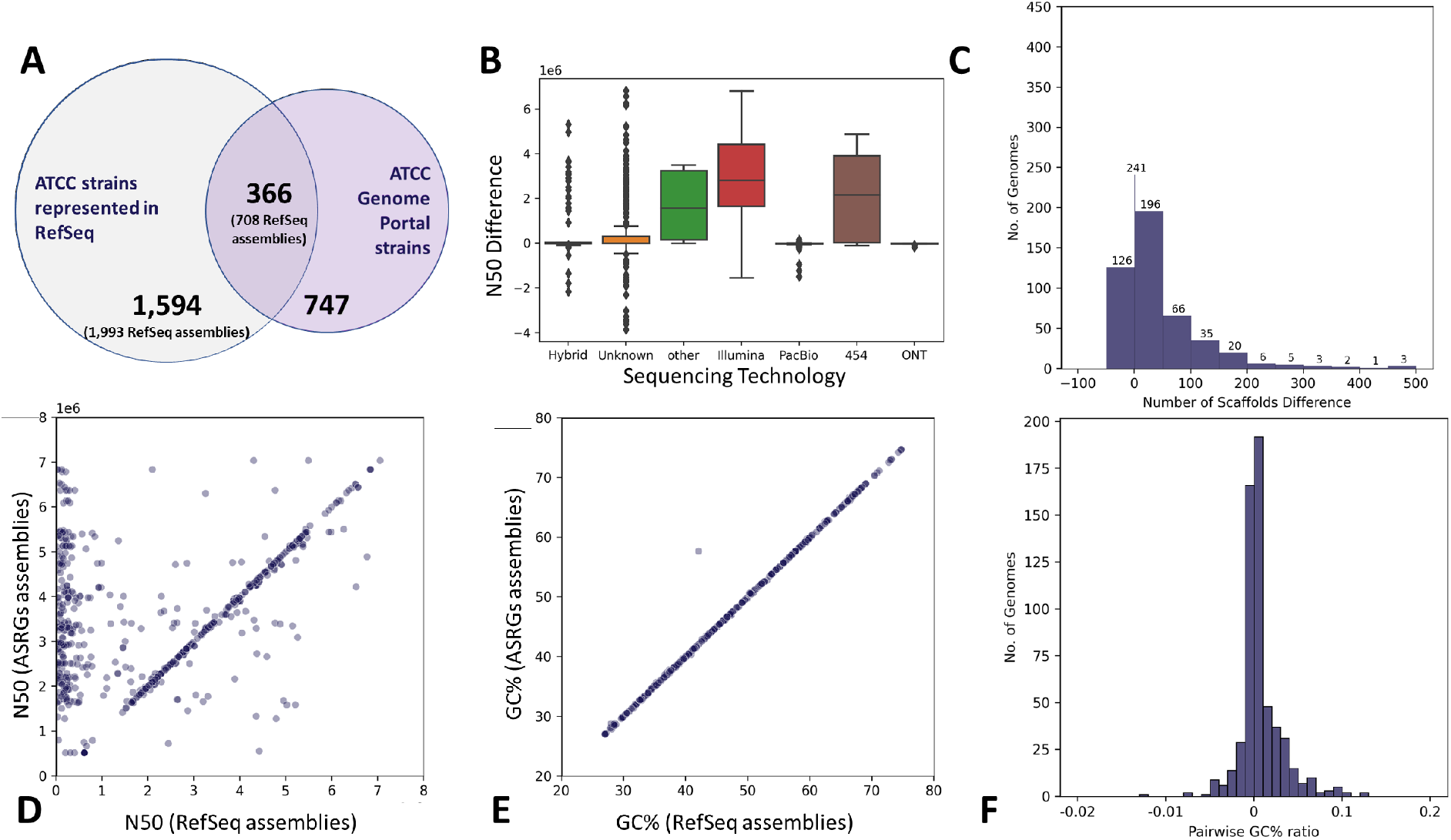
Comparative Metrics for 1,113 ASRGs vs. RefSeq Assemblies. **(A)** Intersection of ASRGs vs. RefSeq for strains labeled as being from ATCC. In parentheses are the total number of RefSeq assemblies, allowing for strain-redundancy. (**B)** N50 variability of RefSeq vs. ASRGs by sequencing technology. Note the scale is 1E6. (**C)** Differences in contig counts for ASRG vs. RefSeq assemblies. Positive values indicating the RefSeq assembly had more contigs. (**D)** Ratios of ASRG N50 values (y-axis) to RefSeq N50 values (“public,” x-axis). Density along the diagonal indicates many assemblies are similar, while the density along the y-axis indicates ASRGs with higher N50 value. (**E)** GC%-content for ASRGs (y-axis) to RefSeq (x-axis). Nearly all assemblies have less than 0.1% difference in GC-content. (**F)** Pairwise GC% differences between ASRGs and comparable RefSeq assemblies for the same strain.

Further examination of the metadata for the 2,701 RefSeq assemblies labeled as ATCC strains also revealed numerous records with incomplete, missing, or obscured descriptor fields (Figure S1). For example, “Assembly type” is present in every assembly record but the value is “na” for all. “Sequencing technology” is not included or has a value of “Unknown” for 1,088 assemblies (∼40%, Table S2), and spelling and nonstandard abbreviations further complicate the rest. **“** Assembly method” is not included for 1,082 assemblies, contains the value “Unknown**”** for 88 assemblies or “other” for 4 assemblies, and has numerous misspellings for various bioinformatics tools (i.e., “Velevt” or “Velveth” for the Velvet assembler). One particularly poor example includes an assembly for *Streptomyces clavuligerus* ATCC^®^ 27064™ (GCF_015708605.1) that indicates the “Assembly method” as “Several assembly pipelines, manual curation v. 2018-09-27.” Underutilized fields included “Description,” “Isolate,” and “Relation to type material,” which had no values in 99%, 98%, and 38% of the assembly records, respectively. The damaging impact that inconsistent depositor metadata has on scientific research and reproducibility has been extensively covered elsewhere ^1,3,25^.

Of the 2,701 RefSeq assemblies for ATCC bacterial strains, 708 had a counterpart ASRG (Table S2, Figure 3A). Of these, 303 (43%) are labeled “complete genome” or “chromosome” level assemblies. Despite this, N50 values were largely inferior when compared to their ASRG counterparts (Figure 3B). While 241 RefSeq assemblies had the same number of scaffolds as their corresponding ASRGs, 341 were more fragmented. Altogether, 662 ASRGs had equivalent or superior N50 values to their RefSeq counterparts (ATCC N50 / RefSeq N50 ≥ 0.95), while 46 ASRG assemblies were more fragmented (Figure 3D). The greatest difference was observed for a RefSeq assembly for *Pseudomonas aeruginosa* ATCC^®^ 700888™ (GCF_000297315.1), which comprised 600 contigs while the ASRG equivalent is closed, containing only one contig.

## Comparative Genomics of 303 RefSeq Assemblies

Next, we compared the 303 complete RefSeq assemblies to their corresponding ASRGs for the same strains (represented by 212 ASRGs). First, we found that the pairwise average nucleotide identity (ANI) ranged from 97% to 100% for identical strains, which at first glance suggested a high level of similarity ^26^. Although large differences in the high-level assembly metrics were previously observed (e.g., N50, GC content), after conducting pairwise whole-genome alignments with *Mummer4* for all 303 RefSeq assemblies against ASRGs for the same strain, we found 292 had over 95% of their sequence aligned. Next, we examined pairwise structural variations and found significant differences in sequence repeats, inversions, indels, and translocations between RefSeq assemblies and ASRGs for the same strains (Tables S3, S4)^27^. Analysis with *dnaDiff* of all 303 RefSeq assemblies revealed an average 6.73 structural rearrangements in comparison to ASRGs, the worst of which was GCF_000160895.1 for *Bacillus cereus* ATCC^®^ 10876™ with 232 structural differences (despite both assemblies having over 99% reciprocally aligned bases). Structural relocations were the most common, with 256 RefSeq assemblies having at least one per assembly (average 4.3 per assembly). Structural inversions were found in 74 RefSeq assemblies (average 2.2). Translocations were relatively rare, with only 9 RefSeq assemblies having structural translocations relative to the ASRG assembly for the same strain (Table S4). We also found that RefSeq assemblies with the greatest number of structural differences from the ATCC assemblies corresponded to those submitted to NCBI prior to 2010, and for which sequencing technology or assembly method were not indicated in the RefSeq metadata. The distribution of structural variations in the 303 complete RefSeq assemblies compared to their corresponding ASRGs is shown in Figure S2.

## Variants in 303 RefSeq Assemblies

Next, we sought to investigate the prevalence of single-nucleotide polymorphisms (SNPs) and insertions/deletions (InDels) that would arise by using RefSeq assemblies as a reference genome against which Illumina sequencing data would be mapped—a common approach used by labs without the resources or expertise for *de novo* assembly and annotation. For each of the 303 complete RefSeq assemblies described above, we mapped the same Illumina reads used in creating the corresponding ASRGs for the same strain. Variant calling from the resulting consensus genomes was carried out on all 303 references to detect SNPs and InDels in each (see Materials & Methods). Overall, the number of SNPs and InDels per assembly ranged from zero (none detected) to as many as 60,064 SNPs *(Acinetobacter baumannii* ATCC^®^ 17978™, GCF_011067065.1) and 2,699 InDels for a given assembly (*Parabacteroides distasonis* ATCC^®^ 8503™, GCF_900683725.1) (Table S5). The median level of SNPs and InDels was 7 SNPs and 8 InDels per assembly, with 7 of the 303 mappings having no detectable SNPs and InDels. These results were promising overall, yet significant outliers were detected, and 26 strains had SNPs and InDels beyond an extreme-outlier boundary, i.e., greater than 3-times interquartile range (IQL) above the median with 9 of them having over 1,000 SNPs and InDels each (Figs. S3, S4a, S4b).

A total of 111 assemblies had fewer than 10 variants, while 15 assemblies had more than 500 variants (SNPs, Indels). Not surprisingly, as the number of SNPs increased, so too did the number of InDels (Figure S3). Of these, 52 of the 303 assemblies had no expected non-synonymous mutations, but 87 had at least 10 non-synonymous variants per genome (Figure S4b). Importantly, 52 RefSeq assemblies identified as “assembled from type material” were found to have at least 10 non-synonymous variants, and seven assemblies had over 100; this could have potentially deleterious impacts on future comparative genomics studies utilizing those reference assemblies (Table S5).

We found that complete RefSeq assemblies without the label “reference genome” or “representative genome” (250 genomes) were enriched for SNPs (7.6-fold) and InDels (9.6-fold) compared to reference RefSeq genomes (53 assemblies). Furthermore, type strain assemblies in RefSeq (i.e., labeled as “assembly designated as neotype,” “assembly from synonym type material,” or “assembly from type material”) had marginally fewer SNPs and InDels than other assemblies overall, but some significant exceptions to this were also observed (see above). No statistically significant enrichment for SNPs or InDels was detectable by taxonomic clade or G:C content. Collectively, these results underscore the importance of data provenance of the originating materials (e.g. “type-strains”) and assembly quality (e.g. “reference genome” or “representative genome”), and that they are both important drivers in reducing variability and improving genome assembly quality.

## Discussion

Over the last 20 years, several non-commercial and government initiatives have specifically tried to address issues relating to the quality and standardization of metadata for microbial genomics, which has had some benefit for end-users, but substantial work remains to be done ^15,28,29^. As the unmet need for curated, high-quality microbial genomics data continues to grow, we will no doubt continue to see a variety of commercial initiatives be successful in developing solutions designed to address gaps in quality, content, and reliability, such as QIAGEN’s CLC Microbial Reference Database, ARES Genetics’ ARESdb, and the One Codex platform. While these public and private efforts have been largely successful, by some measures the overall quality of public microbial genomics data has been declining over the last decade, carrying a potentially great cost to the broader research community ^2,3,5,13,30^. We propose that widespread gaps in the traceability of genome assemblies to their originating biological materials, lab protocols, and bioinformatics methods represent fundamental weaknesses in these data that will hinder research and increase costs unless it is addressed.

At the outset of the work described here, we sought to develop methods to systematically sequence ATCC’s bacterial collection and share that data with the research community alongside the physical strain materials. However, during the course of our work we found that bacterial genome assemblies in RefSeq labeled as representing ATCC strains compared poorly against ASRGs. More broadly, our analysis uncovered disparities in the quality, accuracy, and completeness of metadata associated with assemblies in RefSeq, suggesting that gaps in data provenance may be playing a role in the decline of data quality. As an example, over 33% (1,087) of the RefSeq assemblies included in our study completely lacked any description for how they were sequenced or assembled.

There remain significant gaps in the quality of “typical” genome assemblies available from crowd-sourced databases such as RefSeq. Researchers should be cautious about the data they use and avoid blindly ingesting reference genome data without first being curious about the origins of the data and the methods used to produce them. Further studies are needed to better understand the importance of establishing data provenance in genomics data and the impact its absence has on the research of those who use it. It is our hope that initiatives focused on genomic data provenance, such as the ATCC Genome Portal (https://genomes.atcc.org), will serve to highlight the value of establishing higher standards of traceability and accountability for genomics data in the public domain.

## Materials and Methods

### Sample Acquisition and Culture Conditions

All the bacterial cell cultures and genomic DNA used in this study met or exceeded ATCC’s quality standards (https://www.atcc.org/about-us/quality-commitment), underwent extensive phenotypic and genotypic characterization to ensure accurate strain identification, and were extensively tested for contamination before being accepted for use in this study. ATCC is certified by the ANSI National Accreditation Board (ANAB) to meet both ISO 17034:2016 standards as a reference material producer and ISO/IEC 17025:2017 as a testing and calibration reference laboratory. Each bacterial strain included in this study is available from ATCC’s biorepository and was authenticated according to protocols executed in accordance with ATCC’s quality management system (see above). The specific protocols for each strain varied depending on the specific species in question. In general, strain identification and authentication included assessment of colony morphology, gram staining, culture purity, metabolic profiling, antibiotic susceptibility testing (AST), broad-spectrum biochemical reactivity testing, 16S rRNA gene sequencing, ribotyping, matrix-assisted laser desorption/ionization time-of-flight mass spectrometry (e.g., BioMérieux VITEK MS™ system), and whole-genome next-generation sequencing (NGS). Additional details used for culturing, growth conditions, and authentication of each bacterial strain are available online in each bacterial strain’s catalog page at ATCC.org, and by visiting ATCC’s Bacterial Cell Culture portal^31^.

### DNA Templates and Quality Control

To facilitate the successful NGS library preparation for multiple sequencing platforms (long- and short-read sequences), both high-quality and high-quantity input DNA was obtained from authenticated genomic DNA (gDNA) available in ATCC Bacterial Nucleic Acids repository ^32^. ATCC uses several commercially available extraction kits and in-house validated protocols to obtain pure high-molecular-weight DNA depending on the biological characteristics of the organism undergoing extraction. For strains with no preexisting genomic DNA in ATCC’s repository, total high molecular weight genomic DNA (HMW gDNA) was extracted from thawed or resuspended frozen cultures with 10^7^ −10^9^ cells/mL using the QIAGEN Genomic-Tip™ 20/g or 100/g kit and analyzed for purity, concentration and fragment size. HMW-gDNA samples meeting or exceeding the following criteria were subjected to sequencing; median fragment size larger than 20 kb, optical density A260/280 between 1.75 – 2.00, and a final elution concentration over 20ng/µL per extraction.

### Short-Read Next Generation Sequencing

High-quality gDNA from each strain was subjected to whole-genome sequencing using a short-read next generation sequencing (NGS) workflow. Briefly, sequencing libraries from each extraction were prepared using the DNA Prep kit and indexed using DNA/RNA UD indexes (Illumina), and subsequently subjected to paired-end sequencing on either an Illumina MiSeq® or NextSeq 2000® instrument. Sample multiplexing was based on achieving a minimum 100X average depth of coverage for each genome. Base-calling and adapter trimming was initially done using onboard Illumina instrument software and followed by an additional round of trimming and quality-score filtering using *fastp* and *FastQC* ^33,34^. Illumina reads accepted for further use passed the following quality control thresholds: median Q score, all bases > 30, median Q score, per base > 25, ambiguous content (% N bases) < 5%.

### Long-Read Next Generation Sequencing

Long-read sequencing was carried out using the Oxford Nanopore Technologies (ONT) GridION platform. ONT Ligation Sequencing Kit (Oxford Nanopore, UK, SQK-LSK109) sequencing libraries were prepared from the same physical samples of HMW gDNA used for Illumina sequencing above, multiplexed using the ONT Native Barcoding Expansion kit (Oxford Nanopore, UK, EXP-NBD104 or EXP-NBD114), and sequenced using GridION flow cells (Oxford Nanopore, UK, R9.4.1). As with Illumina sequencing, the number of samples multiplexing was based on the estimated genome size of a given organism and sequencing was performed for a minimum of 48 hours per flow cell. Using the most up to date version of *MinKNOW*, reads were base-called, using the high accuracy settings, demultiplexed, and barcode trimmed. Futhermore, ONT sequencing reads were quality trimmed and filtered using *Filtlong* to meet the following minimum acceptance criteria: minimum mean Q score per read > 10, minimum read length > 5000 bp ^35^.

### Assembly of ATCC Standard Reference Genomes

For genome references deposited to the ATCC Genome Portal, genome assembly size was first estimated from raw reads using *MASH*, and this estimate was used to down-sample the Illumina and ONT raw sequencing libraries to a maximum 100x and 40x coverage respectively^36^. These coverage requirements were selected to maximize accuracy for individual consensus base-calls in the final assemblies^20,22^. After down-sampling each sequencing library, a hybrid *de novo* assembly approach was taken using *Unicyler*^23^. Briefly, Illumina libraries were first assembled individually into contigs. The longest contigs in the initial set were then scaffolded with reads from the ONT library. The combined hybrid-assembly was then iteratively polished using both long and short reads from both input libraries, resulting in highly contiguous or closed reference genomes. Sequencing and assembly artifacts of less than 1000 bp that also had significantly different coverage depth (e.g., “chaff” contigs) were removed from the final draft reference^37^. These draft assemblies were subsequently checked using One Codex to confirm the species^19^. Finally, each draft assembly was assessed for completeness and potential contamination with *CheckM* v1.12, which is based on orthologous gene copy numbers present in an assembly^24^. Assemblies which were determined to have a *CheckM* “completeness” score above 95% and a contamination value below 5% were deemed final assemblies. Each final assembly was subsequently annotated using *Prokka* v1.14 for CDS, rRNA, tRNA, signal leader peptides, and non-coding RNA identification^38^. Finally, each complete and annotated genome was deposited into the ATCC Genome Portal and is referred to herein as an ATCC Standard Reference Genome (ASRG)^39^.

### Characterization of Public Genome Assemblies

To gather the public assemblies of ATCC bacterial strains, the “assembly_summary_refseq.txt” file was downloaded from the NCBI Bacterial RefSeq ftp site (https://ftp.ncbi.nlm.nih.gov/genomes/refseq/bacteria/). This file contains accession numbers and metadata, such as “Isolate”, “Assembly Level”, and “Tax ID,” for every assembly in NCBI Bacterial RefSeq. First, this file was filtered to keep all records that contained either the “ATCC” or “NCTC” keyword. This was done because many strains have synonymous ATCC and NCTC IDs, though often only one of the two is present in a record. Of the records containing “ATCC” or “NCTC,” all that included the “ATCC” were kept, but records containing “NCTC” were filtered to keep only those with a synonymous ATCC ID. This final set of records contained the 2,701 public assemblies of ATCC strains. While “assembly_summary_refseq.txt” does contain metadata, the complete set of metadata was collected by downloading the “assembly_report.txt” for each assembly from the NCBI ftp site. Metadata comparisons were performed using the *compare*.*all*.*levels*.*py* script after appending the RefSeq assembly data with a GC content column, calculated by *bbnorm_stats*.*sh*, all of which was paralleled with GNU Parallels^40^. ATCC’s Genome Portal does not distinguish between contigs and scaffolds, which RefSeq defines as contigs that are connected across gaps. For this, all data comparing ASRGs in terms of contiguity uses RefSeq scaffold information.

### Comparisons of NCBI and ATCC Genome Assembly Metrics

For each of the bacterial strains included in the ATCC Genome Portal, we identified and downloaded all 2,701 genome assemblies that had the same name or similar names from NCBI’s Refseq and Genome Assembly databases. For the 303 NCBI assemblies with a finished assembly status of “Complete” or “Chromosome” and representation in ATCC’s Genome Portal, we carried out pairwise whole genome alignments for each NCBI and ASRG using *MUMmer4* and its associated suite of tools for comparative genomics^27^. In some cases, due to duplications in RefSeq and NCBI’s Genome Assembly database, multiple NCBI assemblies were compared against the same ASRG assembly. Following the creation of the alignments, we identified genome-wide variants for each NCBI assembly as compared to the ASRG assembly, including single nucleotide polymorphisms (SNPs), insertions and deletions (InDels), and structural variants (SV). Genome-wide comparisons using *dnaDiff* v1.3 included assembly length, number of contigs, pairwise percent aligned, and N50 values^41^ (SVs_and_ANI.sh). Furthermore, *MUMmer4’s dnadiff* tool was run with default settings using the ASRG assemblies against each NCBI RefSeq assembly, and relocations, translocations, and inversions are reported alongside total and aligned bases^27^. Prior to running *MUMmer4’s dnadiff* tool on these assemblies, each was filtered to remove contigs <1kb in length to prevent short sequences from exaggerating SVs between assemblies. Structural variants included breakpoints, relocations, translocations, and inversions, and summarized as rearrangements.

## Data Availability

ATCC Standard Reference Genomes (ASRGs), metadata, and raw (FASTQ) data are subject to controlled access, but may be used for any non-commercial research-use only purposes by meeting the requirements outlined below. Data can be obtained directly from the ATCC Genome Portal (https://genomes.atcc.org), via our REST-API (access and details available upon request), or via URLs found in https://github.com/ATCC-Bioinformatics/AGP-Raw-Data/blob/main/AGP_Raw-Data-Access.txt. Downloading these data requires a ATCC Web User Profile (https://www.atcc.org/web-profile/create-a-web-profile) and acceptance of ATCC’s Data Use Agreement (https://www.atcc.org/policies/product-use-policies/data-use-agreement). Any commercial use of ATCC genomics data requires express permission of ATCC (please contact for details). MIT Licensed, open-source code for scripts used in this manuscript are available at https://github.com/ATCC-Bioinformatics/Equivalency_Analysis.

## Acknowledgements

We thank One Codex for contributing to the development of the ATCC Genome Portal. We also thank Drs. Raymond Cypess, Mindy Goldsborough, and Rebecca Bradford for critical comments and review prior to submission.

## Author Contributions

Conceptualization: JGL, BB, JB, JLJ

Data curation: DAY, JGL, NPP, PFC, ALR, MAR

Formal Analysis: DAY, NPP, PFC, ALR

Investigation: CT, AEP, JD, SRG, SK, RM, BB

Project administration: BB, JB

Software: DAY, NPP, PFC, ALR, JB

Supervision: BB, JB, JLJ

Visualization: PFC, BB, JLJ

Writing – original draft: DAY, JGL, JLJ

Writing – review & editing: DAY, NPP, PFC, BB, JB, MAR, JLJ

## Funding

The work described herein was financially supported entirely by the American Type Culture Collection.

## Competing Interests

All authors are employees of the American Type Culture Collection, which solely funded the work presented here and provided all the bacterial strain materials needed for the research. No other competing interests are claimed.

## Supplemental Data

All Supplemental Tables are found in a single MS Excel document, with each worksheet labeled accordingly for each table.

## Supplementary Materials

Supplementary Information is available for this paper. Figs. S1 to S4, and Tables S1 to S8 (as separate Excel document).

Correspondence and requests for materials should be addressed to Jonathan L Jacobs.

**Fig. S1.**
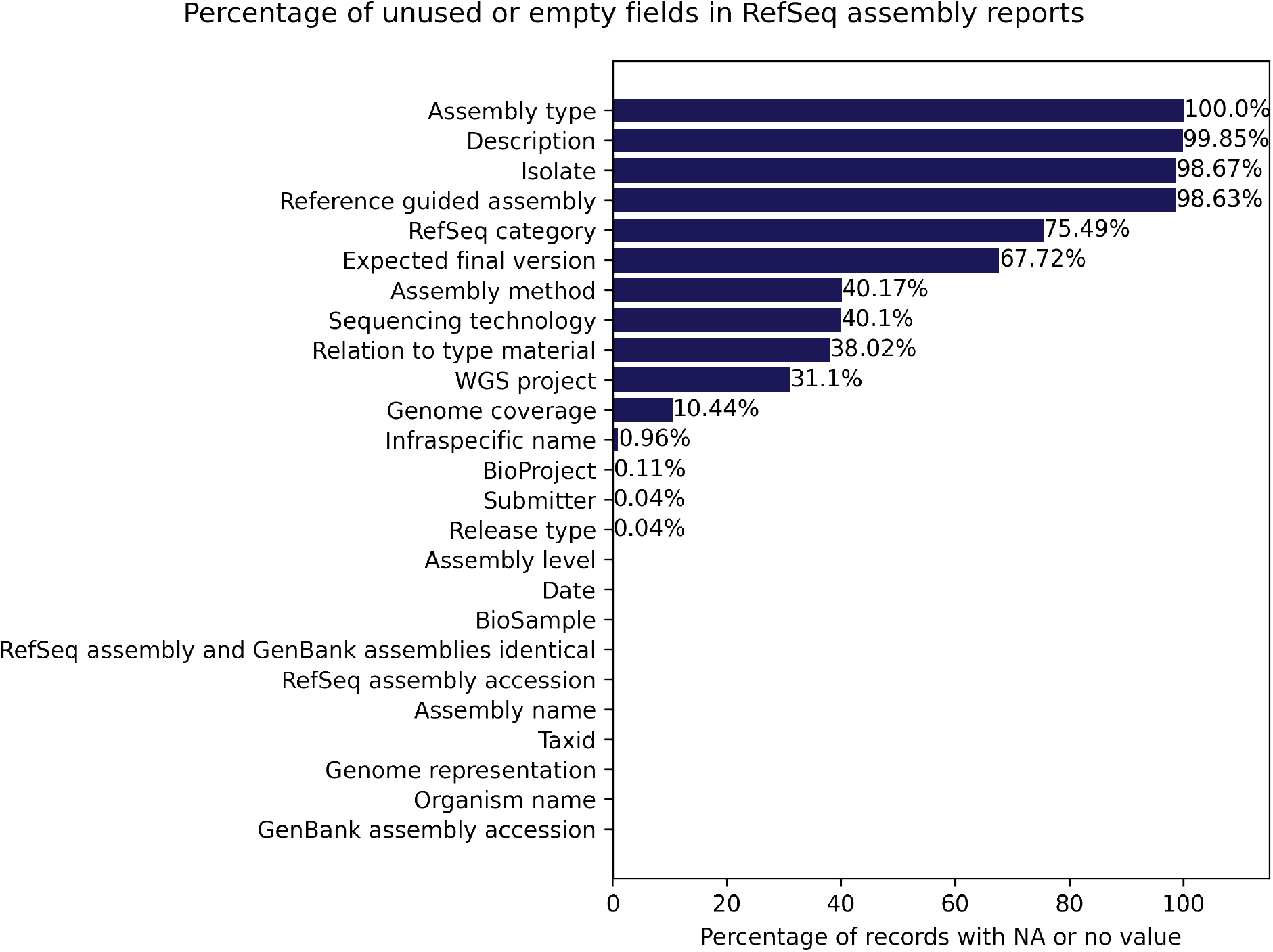
Bar chart demonstrating the percentage of RefSeq assembly report fields that are left empty or contain “na” as a value. While some of these, such as RefSeq category, have implicit definitions for empty fields, others, such as Relation to type material, are potentially crucial pieces of information.

**Fig. S2.**
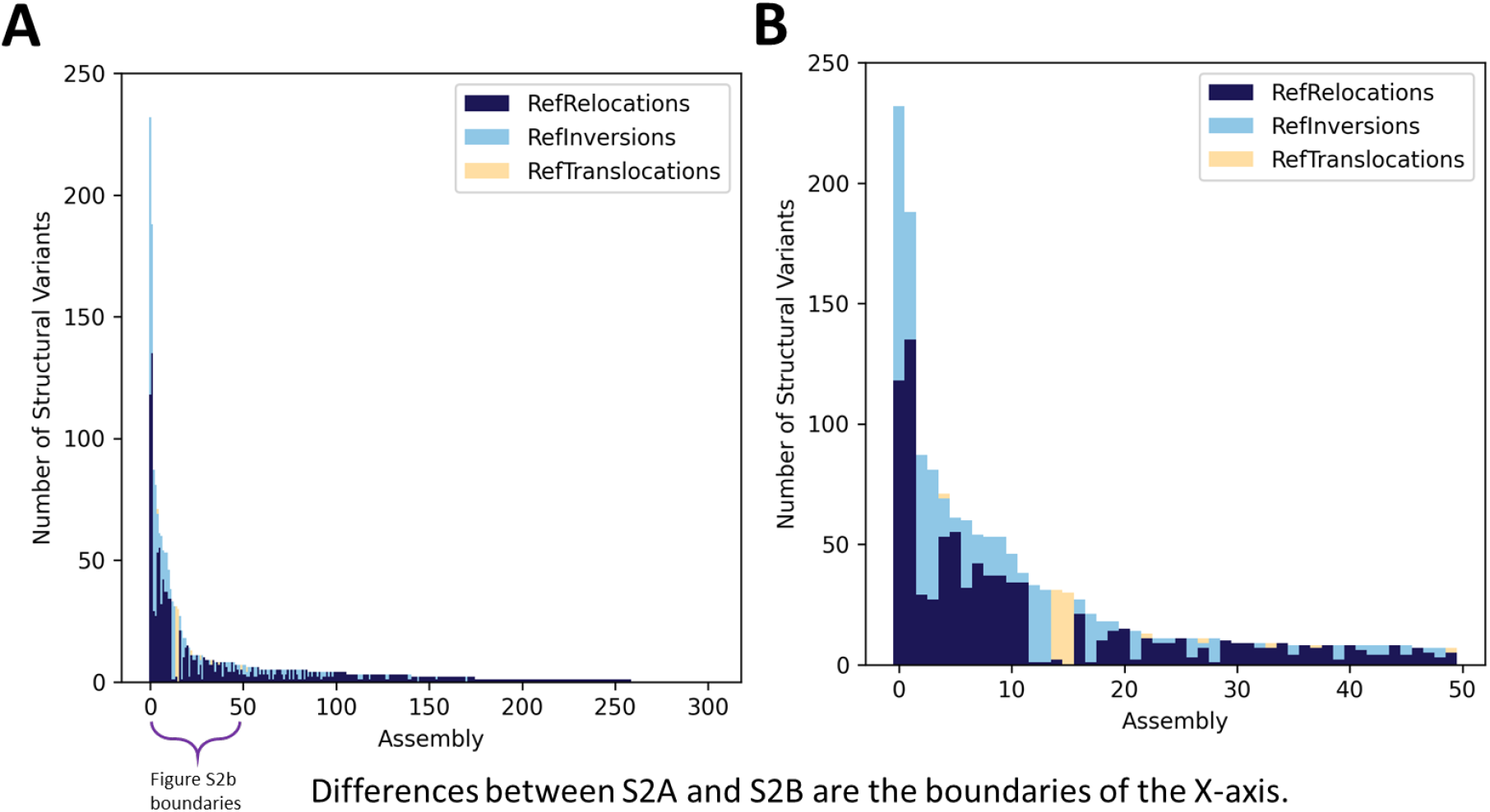
**(A)** Stacked bar chart showing relocation, inversion, and translocation structural variants between all ATCC assemblies and assemblies generated from mapping ATCC’s read data of specific strains to assemblies of those strains. **(B)** Stacked bar chart showing relocation, inversion, and translocation structural variants between ATCC assemblies and assemblies generated from mapping ATCC’s read data of specific strains to assemblies of those strains, for the 50 ATCC products with the greatest total of structural variants.

**Fig. S3.**
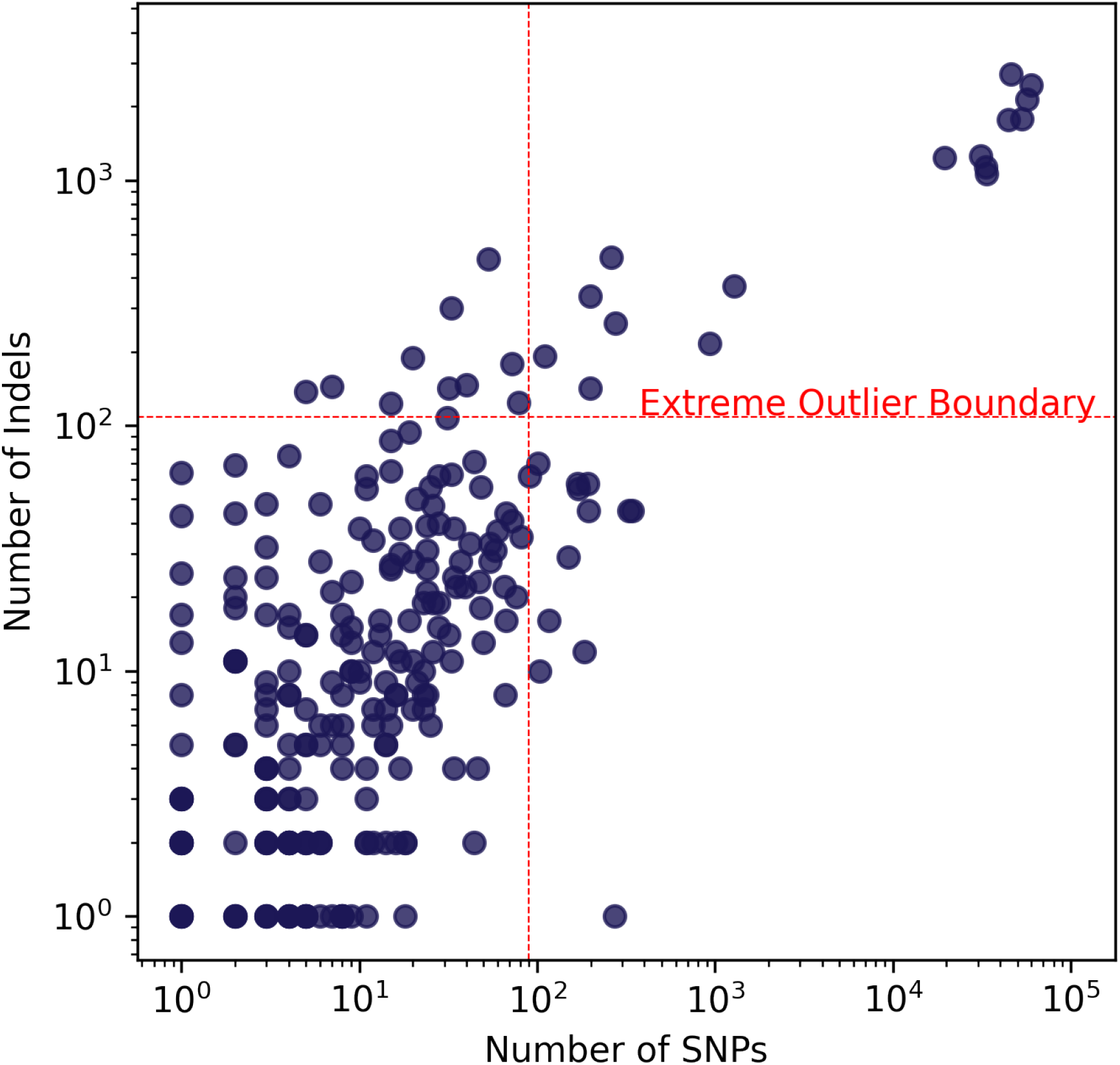
Single-nulceotide polymorphisms (SNPs) and insertions/deletions (InDels) of ASRG raw data mapped to RefSeq references. Each data point represents a read-mapping of ASRG raw data (Illumina only) to a RefSeq genome assembly for the same bacterial strain. In cases where multiple RefSeq assemblies exist for the same bacterial strain, ASRG reads were mapped to each and is represented above by multiple data points. The extreme outlier boundary (red) is determined is 3x the interquartile range above median for both SNPs and Indels (See Materials & Methods).

**Fig. S4.**
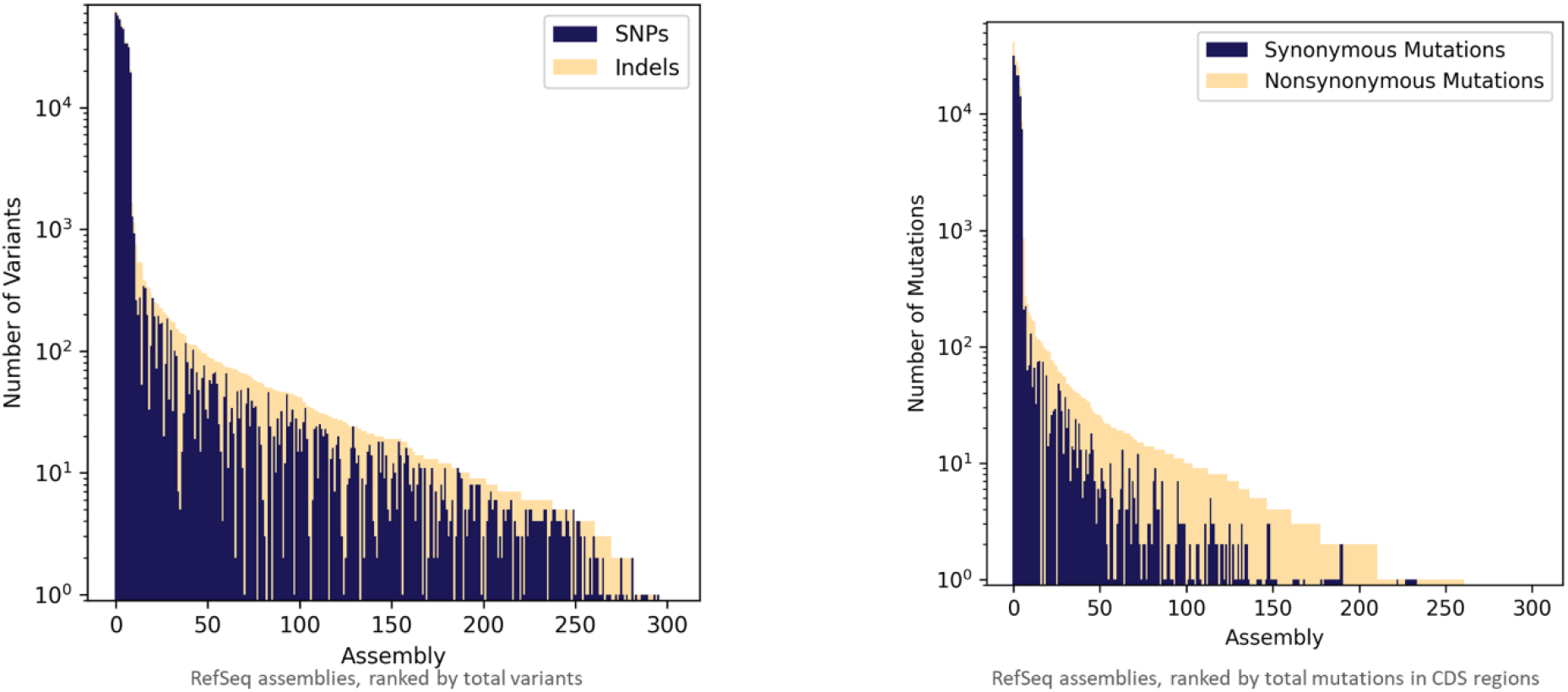
A visualization of the number and types of variants found when mapping the trimmed Illumina reads for an ATCC product to its corresponding RefSeq assembly/assemblies. **(A)** The total number of variants, the number of SNPs, and the number of indels found across this mapping. **(B)** The total number of variants and the characterization of those variants into either Synonymous or Nonsynonymous, as determined by VEP. Synonymous variants represent alterations to a coding sequence that does not change the amino acid upon translation. Nonsynonymous variants represent alterations to a coding sequence that does change the amino acid upon translation. Variants outside of coding regions were calculated as well, but are not shown here.

## Notes

https://genomics.atcc.org

### Competing Interest Statement

The authors have declared no competing interest.

